# A Gene Feature Enumeration Approach for Describing HLA Allele Polymorphism

**DOI:** 10.1101/015222

**Authors:** Steven J. Mack

## Abstract

HLA genotyping via next generation sequencing (NGS) poses challenges for the use of HLA allele names to analyze and discuss sequence polymorphism. NGS will identify many new synonymous and non-coding HLA sequence variants. Allele names identify the types of nucleotide polymorphism that define an allele (non-synonymous, synonymous and non-coding changes), but do not describe how polymorphism is distributed among the individual features (the flanking untranslated regions, exons and introns) of a gene. Further, HLA alleles cannot be named in the absence of antigen-recognition domain (ARD) encoding exons. Here, a system for describing HLA polymorphism in terms of HLA gene features (GFs) is proposed. This system enumerates the unique nucleotide sequences for each GF in an HLA gene, and records these in a GF enumeration notation that allows both more granular dissection of allele-level HLA polymorphism, and the discussion and analysis of GFs in the absence of ARD-encoding exon sequences.

**Abbreviations:**

ARD: Antigen Recognition Domain
EMLB: European Molecular Biology Laboratory
GF: Gene Feature
GFE: Gene Feature Enumeration
HLA: Human Leucocyte Antigen
IHIW: International HLA and Immunogenetics Workshop
IMGT: ImMunoGeneTics
NGS: Next Generation Sequencing
UTR: Untranslated Region

## 1. Introduction

The human leucocyte antigen (HLA) genes are well known as the most polymorphic loci in the human genome. The extensive sequence polymorphism known for the HLA alleles is curated by the ImMunoGeneTics (IMGT)/HLA Database[1], which annotates the individual features for each gene [nucleotide sequences of each exon, intron and flanking untranslated region (UTR)] and gene product (encoded protein sequences). Here, exons, introns and UTRs are collectively referred to as gene features (GFs) to distinguish them from “sequence features” described elsewhere [2].

The World Health Organization Nomenclature Committee for factors of the HLA system (HLA Nomenclature Committee) assigns a unique allele name to each unique HLA nucleotide sequence[3]. Each HLA allele name consists of four colon-delimited fields (e.g., *HLA-A*01:01:01:01*). The first field identifies the allele family (for all genes but *HLA-DPB1*); the second field enumerates the unique protein sequences for the alleles in a given allele family, in the order in which they were identified; the third field enumerates sequences with synonymous substitutions for a given protein sequence, in the order in which they were identified; and the fourth field enumerates sequences with nucleotide substitutions in UTRs and introns for a given synonymous sequence in an exon, in the order in which they were identified. *HLA-DPB1* lacks allele-families; the first field identifies unique protein sequences for all but the *DPB1*02* and *\*04* alleles, for which two distinct protein sequences each are known [3–5].

The IMGT/HLA Database is updated every three months, and the number of named HLA gene and pseudogene sequences increases with each update. For example, 9,946 HLA alleles had been named as of December of 2013[6]; this number increased to 12,242 in December of 2014[7], and 12,542 HLA alleles have been named as of January of 2015. Increases in the number of new allele sequences included in the database have followed the adoption of new genotyping technologies by the Histocompatibility and Immunogenetics (H&I) community, often in conjunction with international HLA and immunogenetics workshops (IHIWs).

The IMGT/HLA Database annotation, based on European Molecular Biology Laboratory (EMBL) formats, is available as hla.dat and hla.xml files from ftp.ebi.ac.uk. These files identify and characterize the nucleotide sequences corresponding to specific GFs for each HLA allele. As illustrated in Table 1, each HLA gene can have a different number of GFs, but all HLA genes have a 3’ and 5’ UTR, at least four exons and at least three introns. However, for most HLA genes, full-length sequence is unavailable for the majority of alleles. As illustrated in Figure 1, nucleotide sequences for more than 60% of HLA-A, -B, -C, and −DRB1 alleles in IMGT/HLA Database version 3.19.0 are available only for exons 2 and 3 of the class I genes, and exon 2 of the class II genes, as these exons encode the antigen recognition domain (ARD). Fewer than 6% of the alleles at these loci have full-length sequences, describing nucleotide sequence for all of an allele’s GFs. Many of these full-length sequences have been generated using next generation sequencing (NGS) technologies, and the number of HLA alleles included in the database seems poised to increase dramatically as NGS technologies become widely used for HLA genotyping by H&I and genomics communities, and as part of the 17^th^ IHIW.

**Figure 1.**
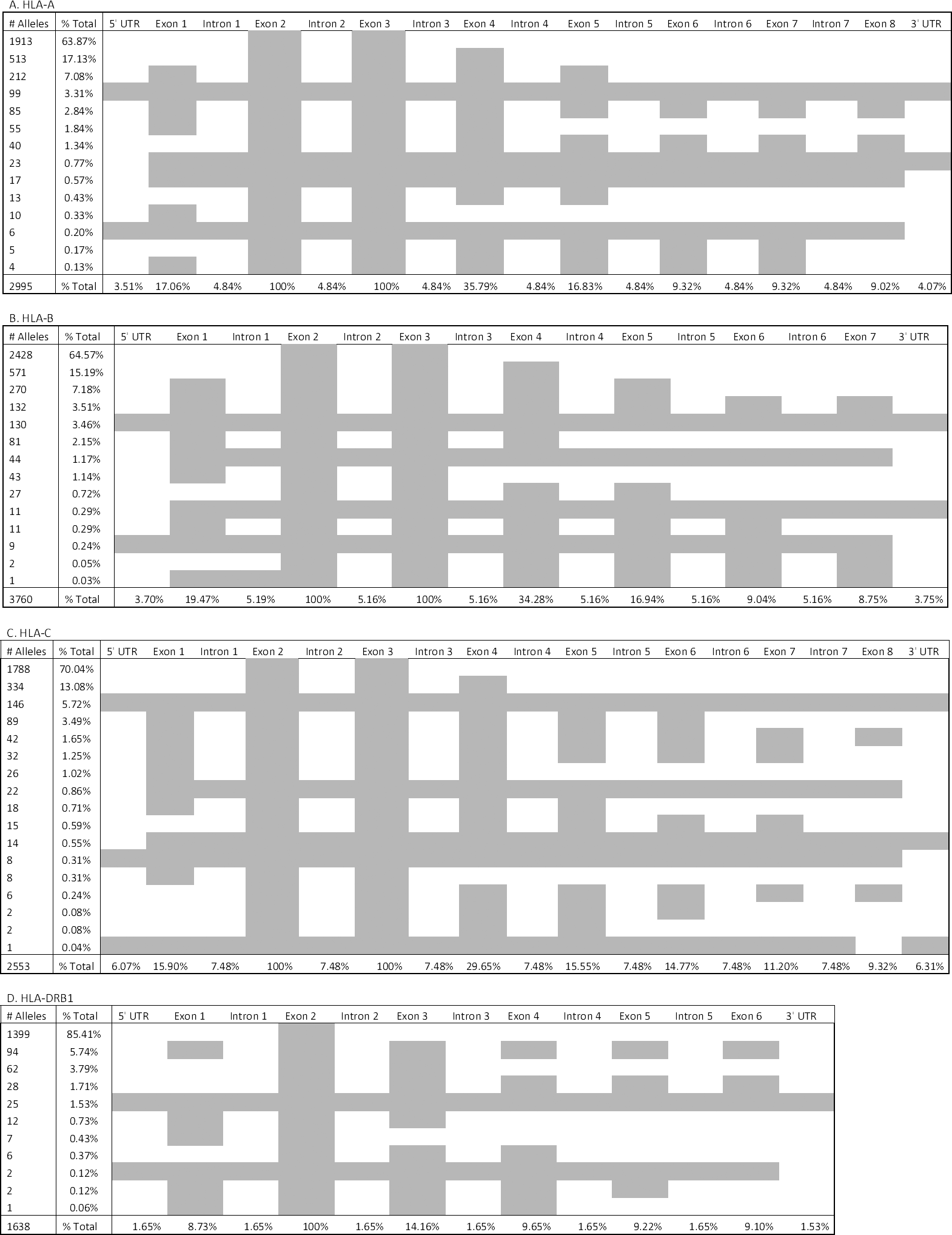
Percentages of *HLA-A, -B, -C* and -*DRB1* Alleles with Nucleotide Sequences for Sets of Gene Features in IMGT/HLA Database Release Version 3.19.0 Each of the four panels details the percentage of alleles for which the nucleotide sequence of sets of gene features (GFs) is known at the *HLA-A, -B, -C*, or -*DRB1* locus. Grey boxes represent GFs for which nucleotide sequence is known for a given percentage of alleles that locus. The % Total value at the bottom of each column represents the percentage of alleles for which nucleotide sequence for each individual GF is known. Each % Total value in the second column represents the percentage of alleles for which nucleotide sequence for the GFs shown in grey in that row are known. The total number of alleles at each locus is shown at the bottom of the first column.

**Table 1.** Maximum Lengths of Gene Features in 11 HLA Genes in IMGT/HLA Database Release 3.19.0

| Locus | 5' UTR | Exon 1 | Intron 1 | Exon 2 | Intron 2 | Exon 3 | Intron 3 | Exon 4 | Intron 4 | Exon 5 | Intron 5 | Exon 6 | Intron 6 | Exon 7 | Intron 7 | Exon 8 | 3' UTR |
| --- | --- | --- | --- | --- | --- | --- | --- | --- | --- | --- | --- | --- | --- | --- | --- | --- | --- |
| HLA-A | 300 | 73 | 130 | 293 | 242 | 314 | 600 | 280 | 102 | 117 | 442 | 33 | 142 | 48 | 169 | 5 | 300 |
| HLA-B | 284 | 73 | 129 | 272 | 250 | 281 | 575 | 277 | 104 | 120 | 441 | 33 | 107 | 44 |  |  | 364 |
| HLA-C | 283 | 73 | 130 | 274 | 250 | 297 | 587 | 277 | 124 | 138 | 440 | 33 | 107 | 48 | 164 | 5 | 171 |
| HLA-DPA1 | 523 | 100 | 3584 | 246 | 340 | 282 | 214 | 155 |  |  |  |  |  |  |  |  | 4331 |
| HLA-DPB1 | 366 | 100 | 4536 | 264 | 4014 | 282 | 547 | 111 | 329 | 20 |  |  |  |  |  |  | 963 |
| HLA-DQA1 | 746 | 82 | 3858 | 249 | 445 | 282 | 429 | 155 |  |  |  |  |  |  |  |  | 436 |
| HLA-DQB1 | 530 | 109 | 1458 | 270 | 2889 | 291 | 517 | 111 | 485 | 24 | 611 | 14 |  |  |  |  | 206 |
| HLA-DRB1 | 607 | 100 | 10306 | 272 | 3464 | 282 | 702 | 111 | 487 | 24 | 1142 | 14 |  |  |  |  | 325 |
| HLA-DRB3 | 327 | 100 | 7681 | 270 | 2302 | 282 | 684 | 111 | 473 | 24 | 799 | 14 |  |  |  |  | 579 |
| HLA-DRB4 | 313 | 100 | 9563 | 270 | 2741 | 282 | 704 | 111 | 474 | 24 | 302 | 14 |  |  |  |  | 570 |
| HLA-DRB5 |  | 100 |  | 270 |  | 282 |  | 111 |  | 24 |  | 14 |  |  |  |  |  |

The maximum length of the nucleotide sequences for each gene feature (GF) [untranslated region (UTR), exon or intron] for each HLA gene in 12,332 HLA alleles in IMGT/HLA Database release 3.19.0 is shown. Blank cells indicate that no GF exists for that gene. Values of 0 indicate that no sequences for that GF have been included in available IMGT/HLA Database annotations.

### 1.1 Application of NGS Technology Highlights Current Nomenclature Limitations

The four colon-delimited field nomenclature for HLA alleles developed in step with genotyping technologies, as greater insights into the nature and scope of HLA polymorphism became available[4, 8–12]. While it provides insight into the *types* of polymorphism that distinguish alleles, this nomenclature does not identify the patterns and location of polymorphism across GFs at a given locus; the extent of the nucleotide sequence represented by an HLA allele name cannot be inferred from that name. The former issue has been partially addressed by extending allele names to identify those alleles that share identical ARD-encoding exon sequences (G groups of alleles, e.g., *HLA-A*01:01:01G*), as well as those alleles that encode identical ARD protein sequences (P groups of alleles, e.g., A*01:01P)[3], as these GFs constitute the largest fraction of the database. However, outside of the G group extension, alleles that share nucleotide sequences for other GFs cannot easily be identified. For example, class I alleles that share identical sequences for one of the ARD-encoding exons, but not the other, cannot be identified using G groups.

The sequences of ARD-encoding exons are required for all nucleotide sequence submissions to the HLA Nomenclature Committee via IMGT/HLA Database, and novel nucleotide sequences for non-coding GFs must be submitted as part of full-length sequences. As a result, an HLA allele name cannot be assigned to a novel nucleotide sequence for an individual GF of interest (e.g., the 3’ UTR of HLA-C [13, 14]) in the absence of nucleotide sequences for ARD-encoding GFs.

Klitz and Hedrick[15] have estimated that millions of alleles persist in the human population for each HLA gene. As NGS technologies extend sequence knowledge into non-ARD encoding GFs, the number of alleles distinguished by synonymous and non-coding variants can be expected to increase dramatically; for example, as illustrated in Table 1, introns 1 and 2 of class II genes can be several thousand nucleotides long, and are likely to have accumulated many nucleotide variants. These variants will be noted in the third and fourth fields of allele names, and it does not seem out of the case to imagine allele names like HLA-DRB1*01:01:100:1004 in the near future. As the number of full-length HLA gene sequences generated increases, it seems likely that a large fraction of them will be unique.

Given the inability to determine which GFs are represented in an HLA allele name, the inability to assign allele names to individual non-ARD-encoding GFs, and the impending likelihood of a large number of unique full-length gene sequences, the utility of the HLA nomenclature is limited for managing, exchanging, discussing and analyzing nucleotide sequences for HLA GFs without the context of ARD-encoding GFs.

Here, a gene feature enumeration (GFE) notation is proposed as a supplement to the current HLA nomenclature for the purposes of cataloging nucleotide sequence polymorphisms for non-ARD-encoding GFs, discussing and analyzing HLA alleles in the context of polymorphism distributed between GFs, and capturing novel nucleotide sequences for non-ARD-encoding GFs generated via NGS technologies. This GFE approach is being developed as part of the 17^th^ IHIWS Informatics Component.

## 2. Gene Feature Enumeration

HLA allele name nomenclature enumerates non-synonymous, synonymous and non-coding nucleotide variants in the second through fourth fields of an allele name. To supplement this approach, the unique sequences in each GF of a given HLA gene can be sequentially numbered, and applied to construct a second name for that allele consisting of one field for each GF, containing the unique number for that GF nucleotide sequence and delimited by colons for consistency with HLA nomenclature, prefaced with the allele name followed by a ‘w’ (for Workshop) to identify the provisional nature of this notation[16, 17]. This GFE notation is illustrated in Table 2.

**Table 2.**
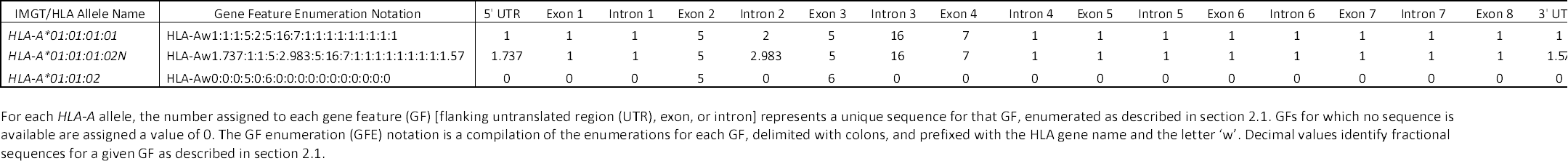
Gene Feature Enumerations for Three *HLA-A* Alleles

For example, the *HLA-A* gene includes 17 GFs (Table 1); any *HLA-A* allele can be represented as a unique haplotype of 17 GFs. As illustrated in Table 2, and in Supplementary Table S1, *HLA-A*01:01:01:01* can be described in GFE notation as HLA-Aw1:1:1:5:2:5:16:7:1:1:1:1:1:1:1:1:1, identifying the constituent sequences of its GFs. In this case, the sequences for the *HLA-A*01:01:01:01* 5’ UTR, exon 1, intron 1, and intron 5 through the 3’ UTR are all the first sequences numbered for those *HLA-A* GFs. The nucleotide sequence of exon 2 is the fifth sequence numbered for that GF, intron 2 the second numbered, exon 3 the fifth numbered, intron 3 the 16^th^ numbered and exon 4 the seventh numbered. The approach applied to assign these GF numbers is described in section 2.1.

In instances where the sequences for all GFs are not known for an allele, a 0 is used to denote unknown sequence. For example, *HLA-A*01:01:02* is one of the 1913 *HLA-A* alleles for which only exon 2 and 3 sequence is known; this allele can be represented as HLA-Aw0:0:0:5:0:6:0:0:0:0:0:0:0:0:0:0:0. By comparing the GFEs for *HLA-A*01:01:01:01* and *\*01:01:02*, it is immediately clear that these alleles share identical exon 2 sequences, and differ only in their exon 3 sequences.

Not all nucleotide sequences for a GF of an HLA gene are the same length. In some cases, these length differences are due to incomplete sequence of the GF in question, and in other cases they are due to insertion-deletion mutations. Using GFE notation, GF nucleotide sequences that exactly match longer nucleotide sequences for that GF are assigned a value equal to the number assigned to the longer sequence plus the fractional length of the shorter sequence relative to the longer sequence. If a 150 nucleotide sequence was an exact match to the first nucleotide sequence of a 300 nucleotide long GF, the 150 nucleotide sequence would be assigned the number 1.5.

For example, *HLA-A*01:01:01:02N* is a null allele that results from a four nucleotide deletion in *HLA-A* intron 2. The GFE for this allele is HLA-Aw1.737:1:1:5:2.983:5:16:7:1:1:1:1:1:1:1:1:1.57. Comparing this GFE to that for *HLA-A*01:01:01:01*, it becomes clear that *the A*01:01:01:02N* 5’ and 3’ UTRs are shorter than those for *A*01:01:01:01* (the 5’ UTR is 73.7% as long, and the 3’ UTR is 57% as long), and that HLA-*A*01:01:01:02N* intron 2 is 98.3% as long as *HLA-A*01:01:01:01* intron 2. *HLA-A* intron 2 is 242 nucleotides long; 238/242 is 0.983.

### 2.1 Assigning Gene Feature Numbers

The GFEs in Figure 2 and Supplementary Table 1 were generated using the hla.xml database export available from ftp.ebi.ac.uk. Nucleotide sequence information for each GF in each HLA gene was isolated and enumerated in decreasing size order, with sequences of the same length ordered by allele name. The longest nucleotide sequence for a given GF was numbered first (to facilitate matching of incomplete sequences), so any GFs with insertion mutations were assigned the lowest numbers.

For example, *HLA-A* exon 2 sequence number 1 was assigned to *HLA-A*23:11N*, which has a 23 nucleotide insertion; sequence 2 was assigned to *HLA-A*68:18N* (20 nucleotide insertion); sequence 3 to *HLA-A*24:232N* (five nucleotide insertion); and 2 sequence 4 to *HLA-A*26:25N* (one nucleotide insertion). Because each of these insertion mutations is unique to one allele, no shorter exon 2 sequences exactly match them, and no *HLA-A* exon 2 nucleotide sequences have been numbered as 1, 2, 3 or 4 plus a fractional length value.

When a nucleotide sequence for a GF is very short, it may exactly match all full length nucleotide sequences for that GF. In these instances, that short nucleotide sequence is numbered as 1 plus the fractional length value. For example, in Table 3, the *DQA1*05:01:02* exon 1 sequence is 13 nucleotides long and is numbered 1.159. The *DQA1*06:01:02* exon 3 sequence is 1 nucleotide long and is numbered 1.004.

**Table 3.** Enumerated Gene Features for HLA-DQA1 in IMGT/HLA Database Release 3.19.0

| IMGT/HLA Allele Name | Gene Feature Enumeration Notation | 5' UTR | Exon 1 | Intron 1 | Exon 2 | Intron 2 | Exon 3 | Intron 3 | Exon 4 | 3' UTR |
| --- | --- | --- | --- | --- | --- | --- | --- | --- | --- | --- |
| HLA-DQA1*01:01:01 | HLA-DQA1w0:1:0:1:0:1:0:1:0 | 0 | 1 | 0 | 1 | 0 | 1 | 0 | 1 | 0 |
| HLA-DQA1*01:01:02 | HLA-DQA1w4:1:12:1:3:1:8:2:1 | 4 | 1 | 12 | 1 | 3 | 1 | 8 | 2 | 1 |
| HLA-DQA1*01:02:01:01 | HLA-DQA1w5:1:19:2:4:1:9:3:2 | 5 | 1 | 19 | 2 | 4 | 1 | 9 | 3 | 2 |
| HLA-DQA1*01:02:01:02 | HLA-DQA1w5:1:13:2:4:1:9:3:2 | 5 | 1 | 13 | 2 | 4 | 1 | 9 | 3 | 2 |
| HLA-DQA1*01:02:01:03 | HLA-DQA1w5:1:14:2:4:1:9:3:2 | 5 | 1 | 14 | 2 | 4 | 1 | 9 | 3 | 2 |
| HLA-DQA1*01:02:01:04 | HLA-DQA1w6:1:10:2:5:1:9:3:3 | 6 | 1 | 10 | 2 | 5 | 1 | 9 | 3 | 3 |
| HLA-DQA1*01:02:02 | HLA-DQA1w0:1:0:2:0:2:0:3:0 | 0 | 1 | 0 | 2 | 0 | 2 | 0 | 3 | 0 |
| HLA-DQA1*01:02:03 | HLA-DQA1w0:1:0:2:0:3:0:3:0 | 0 | 1 | 0 | 2 | 0 | 3 | 0 | 3 | 0 |
| HLA-DQA1*01:02:04 | HLA-DQA1w0:2:0:2:0:2:0:3:0 | 0 | 2 | 0 | 2 | 0 | 2 | 0 | 3 | 0 |
| HLA-DQA1*01:03:01:01 | HLA-DQA1w7:1:15:3:6:4:10:4:4 | 7 | 1 | 15 | 3 | 6 | 4 | 10 | 4 | 4 |
| HLA-DQA1*01:03:01:02 | HLA-DQA1w8:1:11:3:6:4:10:4:4 | 8 | 1 | 11 | 3 | 6 | 4 | 10 | 4 | 4 |
| HLA-DQA1*01:04:01:01 | HLA-DQA1w9:3:20:1:3:1:11:5:5 | 9 | 3 | 20 | 1 | 3 | 1 | 11 | 5 | 5 |
| HLA-DQA1*01:04:01:02 | HLA-DQA1w9:3:16:1:3:1:11:5:5 | 9 | 3 | 16 | 1 | 3 | 1 | 11 | 5 | 5 |
| HLA-DQA1*01:04:02 | HLA-DQA1w0:3:0:1:0:3:0:5:0 | 0 | 3 | 0 | 1 | 0 | 3 | 0 | 5 | 0 |
| HLA-DQA1*01:05:01 | HLA-DQA1w9:4:17:1:3:1:11:1:5 | 9 | 4 | 17 | 1 | 3 | 1 | 11 | 1 | 5 |
| HLA-DQA1*01:05:02 | HLA-DQA1w0:3:0:1:0:1:0:1:0 | 0 | 3 | 0 | 1 | 0 | 1 | 0 | 1 | 0 |
| HLA-DQA1*01:06 | HLA-DQA1w0:0:0:4:0:0:0:0:0 | 0 | 0 | 0 | 4 | 0 | 0 | 0 | 0 | 0 |
| HLA-DQA1*01:07 | HLA-DQA1w8.443:3:16:5:3:1:11:5:5.741 | 8.443 | 3 | 16 | 5 | 3 | 1 | 11 | 5 | 5.741 |
| HLA-DQA1*01:08 | HLA-DQA1w0:0:0:2:0:5:0:0:0 | 0 | 0 | 0 | 2 | 0 | 5 | 0 | 0 | 0 |
| HLA-DQA1*01:09 | HLA-DQA1w0:0:0:2:0:6:0:0:0 | 0 | 0 | 0 | 2 | 0 | 6 | 0 | 0 | 0 |
| HLA-DQA1*01:10 | HLA-DQA1w8.443:1:11:6:6:4:10:4:4.337 | 8.443 | 1 | 11 | 6 | 6 | 4 | 10 | 4 | 4.337 |
| HLA-DQA1*01:11 | HLA-DQA1w5.41:1:18:2:4:1:9:6:2.722 | 5.41 | 1 | 18 | 2 | 4 | 1 | 9 | 6 | 2.722 |
| HLA-DQA1*01:12 | HLA-DQA1w0:0:0:1:0:7:0:0:0 | 0 | 0 | 0 | 1 | 0 | 7 | 0 | 0 | 0 |
| HLA-DQA1*02:01 | HLA-DQA1w1:5:23:8:7:8:12:7:6 | 1 | 5 | 23 | 8 | 7 | 8 | 12 | 7 | 6 |
| HLA-DQA1*03:01:01 | HLA-DQA1w2:5:21:7:1:8:13:8:7 | 2 | 5 | 21 | 7 | 1 | 8 | 13 | 8 | 7 |
| HLA-DQA1*03:02 | HLA-DQA1w3:6:22:7:2:9:13:8:7 | 3 | 6 | 22 | 7 | 2 | 9 | 13 | 8 | 7 |
| HLA-DQA1*03:03:01 | HLA-DQA1w2:5:21:7:1:9:13:8:7 | 2 | 5 | 21 | 7 | 1 | 9 | 13 | 8 | 7 |
| HLA-DQA1*03:03:02 | HLA-DQA1w0:5:0:7:0:10:0:8:0 | 0 | 5 | 0 | 7 | 0 | 10 | 0 | 8 | 0 |
| HLA-DQA1*04:01:01 | HLA-DQA1w0:7:0:9:0:11:0:9:0 | 0 | 7 | 0 | 9 | 0 | 11 | 0 | 9 | 0 |
| HLA-DQA1*04:01:02:01 | HLA-DQA1w12.18:7:4:9:8:12:5:9:10.997 | 12.18 | 7 | 4 | 9 | 8 | 12 | 5 | 9 | 10.997 |
| HLA-DQA1*04:01:02:02 | HLA-DQA1w12.171:7:1:9:8:12:5:9:10.435 | 12.171 | 7 | 1 | 9 | 8 | 12 | 5 | 9 | 10.435 |
| HLA-DQA1*04:02 | HLA-DQA1w12:7:2:9:8:13:5:9:10.243 | 12 | 7 | 2 | 9 | 8 | 13 | 5 | 9 | 10.243 |
| HLA-DQA1*04:03N | HLA-DQA1w0:0:0:10:0:0:0:0:0 | 0 | 0 | 0 | 10 | 0 | 0 | 0 | 0 | 0 |
| HLA-DQA1*04:04 | HLA-DQA1w0:0:0:9:0:14:0:0:0 | 0 | 0 | 0 | 9 | 0 | 14 | 0 | 0 | 0 |
| HLA-DQA1*05:01:01:01 | HLA-DQA1w2.025:8:6:11:9:15:6:9:9 | 2.025 | 8 | 6 | 11 | 9 | 15 | 6 | 9 | 9 |
| HLA-DQA1*05:01:01:02 | HLA-DQA1w10:8:7:11:10:15:6:9:9 | 10 | 8 | 7 | 11 | 10 | 15 | 6 | 9 | 9 |
| HLA-DQA1*05:01:02 | HLA-DQA1w0:1.159:0:15:0:0:0:0:0 | 0 | 1.159 | 0 | 15 | 0 | 0 | 0 | 0 | 0 |
| HLA-DQA1*05:02 | HLA-DQA1w0:0:0:17:0:0:0:0:0 | 0 | 0 | 0 | 17 | 0 | 0 | 0 | 0 | 0 |
| HLA-DQA1*05:03 | HLA-DQA1w10.45:8:8:11:10:16:6:9:9 | 10.45 | 8 | 8 | 11 | 10 | 16 | 6 | 9 | 9 |
| HLA-DQA1*05:04 | HLA-DQA1w0:0:0:12:0:0:0:0:0 | 0 | 0 | 0 | 12 | 0 | 0 | 0 | 0 | 0 |
| HLA-DQA1*05:05:01:01 | HLA-DQA1w11:9:9:11:11:17:2:10:8 | 11 | 9 | 9 | 11 | 11 | 17 | 2 | 10 | 8 |
| HLA-DQA1*05:05:01:02 | HLA-DQA1w11:9:9:11:11:17:1:10:8 | 11 | 9 | 9 | 11 | 11 | 17 | 1 | 10 | 8 |
| HLA-DQA1*05:05:01:03 | HLA-DQA1w11:9:5:11:11:17:4:10:9.554 | 11 | 9 | 5 | 11 | 11 | 17 | 4 | 10 | 9.554 |
| HLA-DQA1*05:06 | HLA-DQA1w0:8:0:11:0:18:0:9:0 | 0 | 8 | 0 | 11 | 0 | 18 | 0 | 9 | 0 |
| HLA-DQA1*05:07 | HLA-DQA1w0:8:0:11:0:16:0:11:0 | 0 | 8 | 0 | 11 | 0 | 16 | 0 | 11 | 0 |
| HLA-DQA1*05:08 | HLA-DQA1w0:9:0:11:0:19:0:10:0 | 0 | 9 | 0 | 11 | 0 | 19 | 0 | 10 | 0 |
| HLA-DQA1*05:09 | HLA-DQA1w0:10:0:11:0:17:0:10:0 | 0 | 10 | 0 | 11 | 0 | 17 | 0 | 10 | 0 |
| HLA-DQA1*05:10 | HLA-DQA1w0:0:0:13:0:17:0:0:0 | 0 | 0 | 0 | 13 | 0 | 17 | 0 | 0 | 0 |
| HLA-DQA1*05:11 | HLA-DQA1w11:9:9:11:11:17:3:12:8 | 11 | 9 | 9 | 11 | 11 | 17 | 3 | 12 | 8 |
| HLA-DQA1*06:01:01 | HLA-DQA1w12.208:7:3:14:12:11:7:9:10 | 12.208 | 7 | 3 | 14 | 12 | 11 | 7 | 9 | 10 |
| HLA-DQA1*06:01:02 | HLA-DQA1w0:0:0:16:0:1.004:0:0:0 | 0 | 0 | 0 | 16 | 0 | 1.004 | 0 | 0 | 0 |
| HLA-DQA1*06:02 | HLA-DQA1w0:0:0:14:0:20:0:0:0 | 0 | 0 | 0 | 14 | 0 | 20 | 0 | 0 | 0 |

### 2.3 Applications of Enumerated Gene Features

GFE notation allows the rapid identification of GFs that are shared by alleles with apparently unrelated allele names. For example, in Table 3, it is clear that several *DQA1*04* and *\*05* alleles (e.g., *\*04:02* and *\*05:01:01:01*) share identical exon 4 sequences; that *\*02:01* and *\*03:01:01* share identical exon 3 sequences, as do *\*04:01:01* and *\*06:01:01*; and that *\*06:01:01* shares identical exon 1 sequence with four *\*04* alleles. In this respect, GFE is similar to the G group approach for identifying alleles with identical ARD-encoding GFs, but allows any combination of GFs to be compared. By identifying the relationship between alleles across all GFs, GFE notation facilitates the investigation of alleles that share, or are distinct at, non-ARD-encoding GFs.

In addition, GFE notation allows alleles that share particular GFs to be combined by converting the enumeration for other GFs to zeros. For example, *DQA1*04:01:01, *04:01:02:01, *04:01:02:02, *04:02* and *\*06:01:01* can be identified as HLA-DQA1w0:7:0:0:0:0:0:0:0, as these alleles all share the same exon 1 nucleotide sequence. In this manner, alleles can be grouped for analysis by the variation at individual GFs, or selected GF sets, without having to parse the sequence information represented by the allele name. In this respect, GFE notation offers a solution to the potential problem of evaluating the significance of hypothetical alleles such as “HLA-DRB1*01:01:100:1004” and “HLA-DRB1*01:03:76:408”; using these allele names, the simple option for analysis would be to truncate them to two-field names (*HLA-DRB1*01:01* and *\*01:03*), ignoring any synonymous or non-coding variation. Using GFE notation, it may be possible to identify GFs that distinguish these alleles in terms of analytical significance.

Finally, GFE allows the consideration of nucleotide sequence variants in non-ARD-encoding GFs, and specifically in the absence of any non-ARD-encoding GF sequences. For example, if 3' UTR sequences are being studied, those 3' UTR sequences can be considered in the context of known polymorphism at a locus by using GFE notation that pertains to the 3' UTR only (e.g., HLA-DQA1w0:0:0:0:0:0:0:0:7 vs HLA-DQA1w0:0:0:0:0:0:0:0:10). GFE notation also allows for novel GF sequence variants to be compared with sequence variants that are already in the IMGT/HLA Database by assigning a new number to the novel variant. For example, the database includes 10 unique *HLA-DQA1* 3’ UTR sequences; a novel *HLA-DQA1* 3’ UTR can be named with GFE notation as HLA-DQA1w0:0:0:0:0:0:0:0:11 in the absence of exon 2 nucleotide sequences.

### 2.3 A Service for Managing Gene Feature Enumeration

GFE notation is not intended to replace HLA allele names; the long histories of HLA nomenclature and the H&I field, and the notoriety of specific HLA allele names preclude these names from ever being retired. GFE notation is proposed as a means for managing and discussing HLA polymorphism by acknowledging the underlying structure of the genes. As such, it is best managed in an automated fashion, with new GFE notations added as new HLA alleles are named, and updated as the nucleotide sequences of extant HLA allele names are extended to new GFs, with each IMGT/HLA Database release. An internet-based service would make GFE notations publically accessible, and would permit the automated inter-conversion of allele names and GFE notations. This service is under development as part of the 17^th^ IHIW Informatics Component (ihiws.org/informatics-of-genomic-data/), and will be made available as an open-source product. Such a service could be applied to other highly polymorphic genetic systems.

Clearly, GF-level knowledge of the elements that distinguish alleles will be insufficient for many use cases. This service would serve a second function of characterizing the nucleotide polymorphisms that distinguish unique sequences for a given GF, fostering more granular investigations of HLA polymorphism.

The final function of this service would be to register novel, uncurated HLA gene sequences generated by 17^th^ IHIW NGS projects, and eventually by any HLA sequencing effort, prior to submission to the IMGT/HLA Database, or for instances when ARD-encoding GF sequences are not available. Undoubtedly, many of the sequences so registered will result from sequencing errors, and should not be included in the IMGT/HLA Database. However, genuine novel nucleotide sequences may presumably be reported by multiple sequencing efforts, and this service would serve as a clearing house for such sequences.

## 3. Conclusions

Gene feature enumeration is a novel approach to describing HLA polymorphism that takes the structural elements of an HLA gene into account. This approach should be relatively easy to adopt, because it relies on information resources already provided by the IMGT/HLA Database. GFE notation will not supplant HLA allele names, but can complement them as the database accumulates sequences for non-ARD GFs generated via NGS methods. As knowledge of the regulatory roles of non-coding nucleotide sequences and their functional impacts on the HLA genes grows, it seems possible that the definition of an HLA allele may someday include promoters, enhancers and other intergenic sequences. GFE enumeration can accommodate this kind of growth in our understanding of immunogenetics in ways that allele names cannot. Given the inevitable changes that NGS methods will have on the H&I field, now is the time to discuss the best means of adapting to them.

## Acknowledgements

This work was supported by National Institutes of Health (NIH) grants U01AI067068, awarded by the National Institute of Allergy and Infectious Disease (NIAID), and R01GM109030, awarded by the National Institute of General Medical Sciences (NIGMS). The content presented is solely the responsibility of the author and does not necessarily represent the official views of the NIH, NIAID, NIGMS or United States Government. The input of Henry Erlich, Marcelo Fernandez-Viña, Damian Goodridge and Martin Maiers is very much appreciated in the development of this work.

